# Using qualitative models to define sustainable management for the commons in data poor conditions

**DOI:** 10.1101/063479

**Authors:** Francesca Mancini, George M. Coghill, David Lusseau

**Author notes:** Corresponding author: Francesca Mancini, Room 418, Zoology Building, Tillydrone Avenue, Aberdeen AB24 2TZ, UK.

## Abstract

Nearly 50 years after Hardin’s “tragedy of the commons” we have not yet found predictive tools to guide us towards sustainable management of common-pool resources (CPR). We often have a good understanding of the qualitative relationships between the principal actors in socioecological systems (SESs), but classical quantitative approaches require a tremendous amount of data to understand the drivers of SESs sustainability. Here we show that qualitative modelling approaches can provide important governance insights for SESs that are understood but not quantified.

We used Loop Analysis to test the outcomes of different management regimes on a simple nature-based tourism SES described by economic, social and environmental variables. We tested the sustainability of different management scenarios on this system and we identified the necessary conditions to achieve it.

We found that management regimes where property rights and responsibilities are shared between different stakeholders are more likely to be successful. However, the system is generally highly unstable and it is important to tune each strategy to each particular situation.

The conditions for sustainability found across the different systems tested were: a low reinvestment rate of tourist revenues into new infrastructures and a low growth rate of the environment. Management strategies based on maximum sustainable yield, which keep the environment far from its carrying capacity, have less chance to be sustainable.

Qualitative models of SESs are powerful diagnostic tools; they can help identifying variables that play an important role in determining socioecological sustainability in data-poor circumstances and guide the design of efficient data collection programmes. Our results highlight the importance of careful planning when designing management strategies for nature-based tourism. The application of one-size-fits-all solutions to every situation is likely to lead to the failure of the commons; however tourism-based SESs can be sustainable if management strategies are tuned to each particular case.

## 1. Introduction

Natural resources are usually considered common-pool resources (CPRs): it is usually impossible or very costly to exclude individuals from using them and their use by one user reduces the quantity or the quality available to other users (Ostrom et al., 1999). There are two main approaches to dealing with the “commons dilemma”. The “panacea” approach applies simplified and general models to all situations. Advocates of this approach propose one particular governance structure as the only possible solution to the tragedy of the commons (Hardin, 1968). The other approach consists in deriving from empirical case studies the characteristics that enable sustainable governance (Ostrom, 1990). The first approach does not recognise the importance of the particular circumstances that characterise each different situation (Ostrom et al., 2007), while the second has to deal with all the issues associated with obtaining observations and data from these complex socioecological systems (SESs)(Hilborn and Ludwig, 1993). As a consequence of the limitations of these approaches, attempts to manage CPRs have often failed (Acheson, 2006).

Commons and their users form SESs, which are composed of different, relatively separable, subsystems that interact in a complex and, sometimes, unknown way (Ostrom, 2009). The inherent complexity of SESs requires an integrated approach to predict the outcomes of management strategies (Ostrom, 2007). However, we do not yet have analytical tools to accurately predict these outcomes (Agrawal, 2014), especially in data-poor circumstances. These systems are difficult to study empirically, because the scope for experimental work is limited and replication, control and randomisation are difficult to achieve (Hilborn and Ludwig, 1993). Therefore, a simulation approach could offer insights on the outcomes of different management regimes. However, little is known about the relationships between the ecological and socio-economic components of these systems and, often, we cannot quantify important variables in the model. Qualitative approaches have proven advantageous to model complex systems in data-poor circumstances (Metcalf et al., 2014).

Recreation is one of the cultural ecosystem services that the environment provides. Tourism is often a primary income for local communities, it can dominate national economies and play a key role in nations’ macroeconomics (O’Connor et al., 2009). While nature-based tourism has been welcomed by conservation and environmental organisations as an eco-friendly alternative to other consumptive activities, such as hunting and fishing (Tisdell and Wilson, 2002), there is growing evidence that nature-based tourism, if not managed properly, can have negative effects on the environment (Meletis and Campbell, 2007; Pirotta and Lusseau, 2015). Therefore, the issue of managing nature-based tourism becomes a CPR issue.

In this study, we tested the sustainability of management regimes on qualitative representations of nature-based tourism SESs using Loop Analysis (Puccia and Levins, 1985). SESs are subjected to press-pulse dynamics (Collins et al., 2011) and in order to understand what drives their sustainability we need to investigate their responses to both press and pulse perturbations. Pulse perturbations are sudden events, such as droughts or fire, which rapidly alter the state of the system, while press perturbations are sustained and slow, such as climate change or economic growth. A pulse event temporarily “shakes” the system, while a press disturbance slowly pushes it away from its current state. We define sustainability in terms of responses of the SES to pulse and press perturbations. For each different management strategy applied to a simple nature-based tourism system we asked three questions: 1) Does the system’s equilibrium lose stability after a pulse perturbation? Stability is the ability of a system to return to its previous state after a perturbation. A stable system offers more predictability and reliability of management interventions, because it is less likely to shift to a different state after a sudden event. We assessed this property of the system using qualitative stability criteria. 2) Under which conditions could the system remain stable? A sensitivity analysis of the stability criteria can identify the key drivers of system’s stability, in other words, which components of the system could be modified to shift the system from being dysfunctional and unstable to being functional and reliable. 3) How does the system behave after a press perturbation? For example, during the development of a nature-based tourism destination, how will the different components of this system respond to an increase in the number of tourists using the area? If this positive press perturbation does not result in environmental degradation, or a reduction in the number of users or in the tourism capital, then the SES can maintain environmental quality, social justice and economic profitability, in other words, triple bottom line (TBL) sustainability (Elkington, 1998). In this study, social justice is intended as access to the resource by the community of users: if responses to press perturbations predict that some users will be excluded from the resource we considered the system to be unjust. For a system to be sustainable it needs to be stable to pulse perturbations and have potential to keep TBL sustainability in presence of a press disturbance in any of its components.

## 2. Materials and Methods

### 2.1 Property rights scenarios

In the resource management literature property is mainly considered as owned or affected by private individuals, local communities or governments (Acheson, 2006; Hoffmann, 2013). In this study we consider an open access scenario in which there are no rules governing property rights, and scenarios where property rights are owned by a central authority or the local community of users. In order to represent both marine and terrestrial systems, we do not consider private property, which is often not possible in a marine context where boundaries are difficult to define and wildlife is highly mobile. Some studies have highlighted the importance of nesting and institutional variety in governance structures (Dietz et al., 2003), showing how mixed strategies can determine the success of CPRs (Pirotta and Lusseau, 2015). Following these studies we considered hybrid scenarios, where property rights are shared between the users and a third party. Within these property rights regimes we also considered different management tools.

The scenarios are represented as signed digraphs (Fig. 1). The nodes represent the variables in the system. The links connecting the nodes represent the qualitative relationships between the variables. Positive relationships (an increase in the first variable produces an increase in the abundance of the second variable) are represented by links with an arrow-end, while the links with a circle-end represent negative relationships. Links that start and end on the same variable are called self-effects, and they represent self-regulation (e.g. density dependence) or reliance on factors external to the modelled system.

**Fig. 1.**
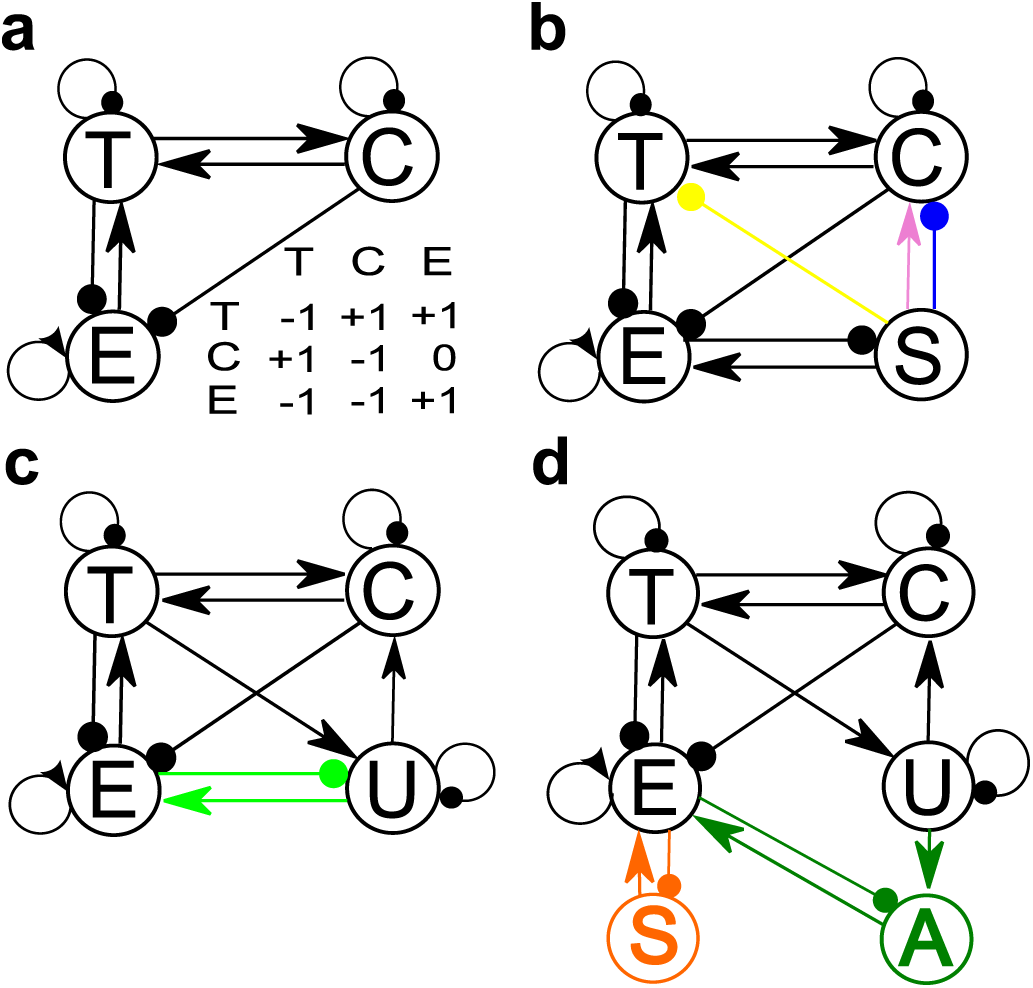
Signed digraph of all the scenarios tested. The nodes represent the variables in the model. T: tourists; C: capital; E: environment; S: state intervention; U: users; A: external agency. The links connecting the nodes represent the relationships between the variables: arrow-ended links indicate positive relationships, circle-ended links represent negative relationships. The links starting and terminating on the same variable represent self-effects. a) Signed digraph of the open access scenario and its matrix representation. Each entry in the matrix corresponds to a link in the graph. b) State ownership scenarios. The pink (dashed), blue (dash-dotted) and yellow (dotted) links represent the three alternative scenarios, respectively, subsidies, licencing and access fee. c) User group ownership scenarios. The first scenario is represented by the black links, while the second scenario includes the green dotted links representing the adaptive management of the environment. d) Hybrid scenarios. In the first scenario the government intervenes to monitor and manage environmental quality (orange dotted lines), in the second one users invest in an external agency to monitor and manage environmental quality (green dashed lines). For detailed description of the models see text.

#### 2.1.1 Open access

This scenario (Fig. 1a) describes an unregulated system where access and use of the resource are unregulated. This core model builds on previous work on sustainable tourism by (Casagrandi and Rinaldi, 2002); the equations presented by Casagrandi and Rinaldi were converted into a signed digraph and all the following scenarios are derived from this core model by adding feedbacks representing governance structures. This simple model represents all main system components: tourists (T), the capital (C), intended as structures available for tourism activities, and the environment (E). The users of the resource are the tour operators that offer, for example, wildlife watching trips or guided walks. In this model the resource users are included in the variable C. All the links to the variable T (Fig. 1) represent the attractiveness of the site; tourists are attracted by the presence of amenities and environmental quality, while the attractiveness of the site decreases with overcrowding. The infrastructure degrades and investment is needed to renew it. Tourists generate revenues that are invested in new infrastructures. Tourism infrastructures, such as hotels, vehicles for wildlife or sightseeing tours, roads etc., negatively impact the environment as do the tourists (Casagrandi and Rinaldi, 2002). The environment is assumed to have a carrying capacity with density-dependence effects (Casagrandi and Rinaldi, 2002) and it is not assumed to be in pristine conditions in absence of tourism. Therefore the environment exploited by human activities is kept far from its carrying capacity and will exhibit a positive self-regulation. We do not assign magnitudes to the effects just described and, therefore, they can range from negligible, to very strong ones.

#### 2.1.2 State ownership

We built different state-ownership property rights scenarios by adding a variable for state intervention (S) to the open access model (Fig. 1b). We developed three scenarios. State intervention is stimulated by a reduction in environmental quality in all the models and the state always implements measures to improve environmental quality. This negative loop between S and E represents an adaptive management strategy. The subsidies scenario (pink dashed line in Fig. 1b) represents a situation in which the state subsidises the industry to build new infrastructures. In the licensing scenario (blue dash-dotted line in Fig. 1b), the state holds property rights on the resource and limits the expansion of the infrastructure by issuing licences to a restricted number of users. In the access fee scenario (Fig. 1b, yellow dotted line), state intervention controls the number of tourists allowed in the area (e.g., entrance fee for a national park).

#### 2.1.3 User group ownership

In these two scenarios (Fig. 1c) property rights are owned by the users’ group, which becomes an explicit variable in the model, U. Users have the right to access, use and manage the resource, exclude other individuals from the resource and have alienation rights. The positive links from T to U and from U to C represent this alienation right; according to the flow of tourists, the users can decide to retain or sell their right to access, use and manage the resource, thus increasing or decreasing the number of structures available to the tourists (for example the number of boats available for trips). In the second scenario the users also implement an adaptive management strategy, represented by the negative feedback loop between E and U (green dotted lines in Fig. 1c): a decrease in environmental quality stimulates an intervention from the users that will improve the health of the environment.

#### 2.1.4 Hybrid

In the hybrid scenarios (Fig. 1d) property rights are shared between the users’ group and a third party. The users still have access, use, exclusion and alienation rights, but the government (Fig. 1d, orange dotted lines) or an agency funded by the users (Fig. 1d, green dashed lines) is in charge of monitoring and managing environmental quality. The first model represents a situation where the government is required to monitor the environment, often because of international agreements on environmental quality. In the second model, the tour operators fund the environmental monitoring and management themselves, for example by paying an environmental agency.

### 2.2 Loop Analysis

Here we briefly describe the theory of Loop Analysis (Puccia and Levins, 1985). Model construction can start with the signed digraph (Fig.1). In the graph, a path is defined as a series of links starting at one variable and ending on another without crossing any variables twice while a set of links that starts and ends at the same variable is called a loop. Loops in the same system are defined as either conjunct or disjunct: two conjunct loops have at least one variable in common (in Fig.1a the self-loop of E and the loop between E and T), while two disjunct loops have no variables in common (in Fig.1a the self-loop of E and the loop between T and C). The qualitative relationships represented in the graph can be entered into a matrix:

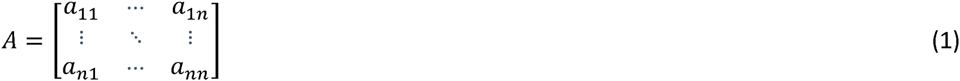

where element *a*_*ij*_ represents the effect of variable *j* on variable *i*. The matrix represents qualitative relationships (negative, positive or 0), without specifying the magnitude of the effects (Fig.1a).

#### 2.2.1 Stability to pulse perturbations

Stability of loop models is assessed by examining the system’s feedback. A feedback loop can be positive or negative according to the product of its links. The loop between T and C in Fig. 1a is positive (a_CT_ * a_TC_= (+1) * (+1) = +1) while the one between T and E is negative (a_TE_ * a_ET_ = (+1) * (−1) = −1). A system adequately regulated by negative feedback is stable because after any perturbation, negative feedback will bring the system back to its original state. A system has multiple feedback paths of different lengths, called feedback levels: within an ecological system there will be feedback paths of length two, such as predator-prey relationships, and feedbacks of length three, such as two predators competing for the same prey. The feedback levels of a system range from 1 to the total number of variables and feedback at each level *k* is calculated as the sum of all the loops of length *k* plus the products of disjunct loops that have combined length *k*.

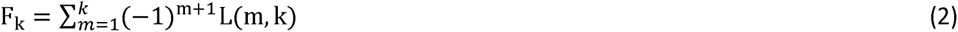

The notation L(m, k) means the product of *m* disjunct loops of combined length *k*. For example L(2, 4) means the product of two disjunct loops of combined length 4 (the product of two disjunct length-two loops or one self-loop multiplied by one length-three loop). *m* goes from 1 to a maximum of *k* because feedback at higher levels is composed of a combination of feedbacks at lower levels plus longer pathways and the number of disjunct loops *m* cannot exceed k. If *k* = 3, *m* can go from 1 to 3 because the system can have one loop of length three, two disjunct loops of combined length three, such as a self-loop and a length-two loop, and three self-loops of combined length three, but it cannot have 4 disjunct loops of combined length three. The term (−1)^(m+1)^ is a sign adjusting factor that tells us that if the number of disjunct loops (*m*) multiplied together is even, then their product is multiplied by −1. This is to avoid that an even number of negative loops multiplied together give a positive feedback term.

The first feedback level for the system described in Fig. 1a is given by the sum of all the self-loops:

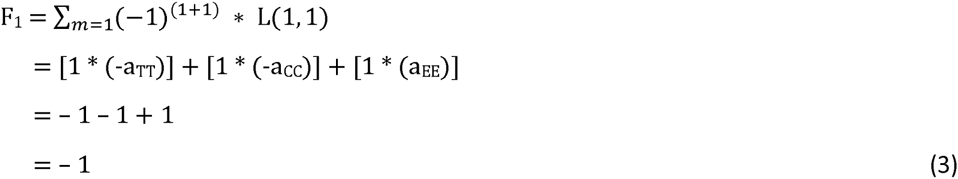

The second feedback level for the system described in Fig. 1a is given by the sum of all the length-two loops (the loops between T and C and between T and E) plus the products of all the pairwise combinations of self-loops:

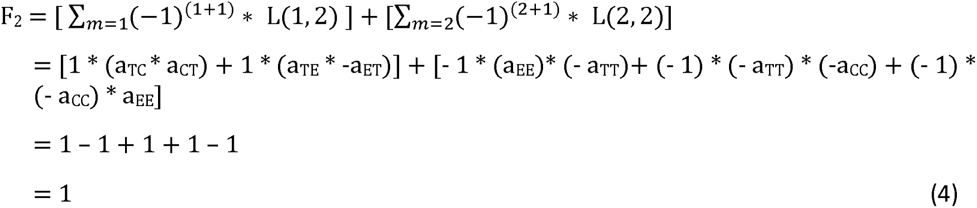

The third and last feedback level is given by the length-three loop (between E, T and C), plus the products of all the combinations of disjunct length-one and length-two loops (for example the self-loop of C and the loop between E and T) plus the product of the three self-loops:

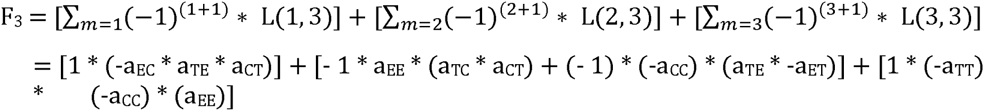

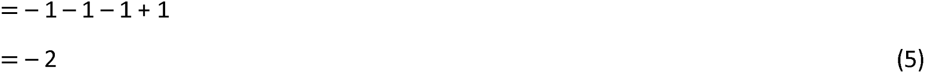

Loop Analysis considers two criteria for stability, the Routh-Hurwitz criteria (Hurwitz, 1895; Routh, 1877). The first criterion states that feedback at all levels is negative (Routh, 1877). According to this criterion the system described in Fig. 1a is unstable because F_2_ is positive (Eq. 4). The second criterion depends on feedback at lower levels of the system being stronger than feedback at higher levels. This is because a system dominated by higher-level feedback tends to overcorrect and any disturbance will be amplified through increasing oscillations. This second criterion is satisfied by the Hurwitz determinants (Hurwitz, 1895) being positive:

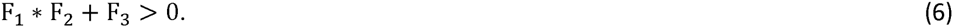

Eq. 6 defines the second Hurwitz determinant, which is used as second condition for stability for loop models with three and four variables (Puccia and Levins, 1985). For the system in Fig. 1a the third Hurwitz determinant is (−1) * 1 + (− 2) = − 3. Hence, the open access system in Fig. 1a is unstable according to the both stability criteria. For loop models with five variables the formula for the third Hurwitz determinant applies:

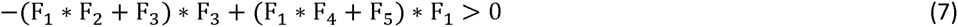

It is also possible to conduct a sensitivity analysis of each feedback level and Hurwitz determinant to each element in the matrix (Hosack et al., 2009). In other words, which direct effects in the system are crucial to obtain a negative feedback and a positive Hurwitz determinant? The procedure counts how many times each direct effect in the model appears in stabilising (negative for feedback levels, positive for Hurwitz determinants) and destabilising (positive for feedback levels, negative for Hurwitz determinants) elements (feedback cycles) in the calculation of Hurwitz determinants or feedback levels and divides it by the total number of feedback cycles in which the same direct effect appears. The index takes values from −1 to 1; for sensitivities of feedback levels, values close to −1 indicate that the direct effect appears only in stabilising feedback cycles, a value of 0 indicates that the direct effect appears in the same number of stabilising and destabilising feedback cycles, while a value close to +1 indicates that the direct effect has a highly destabilising effect on the system. The opposite is true for sensitivities of Hurwitz determinants. For the feedback level calculated in Eq. 5, the direct effect a_EE_ appears in two feedback cycles, one negative or stabilising (− 1 * (a_EE_) * (a_TC_ * a_CT_)) and one positive or destabilising (1 * (−a_TT_) * (−a_CC_) * (a_EE_)). Therefore, the sensitivity index of the feedback at level three to the self-loop of E is: (1 – 1)/2 = 0. For a detailed description of this method see (Hosack et al., 2009).

For each of the systems described in section 2.1 we used the Routh-Hurwitz criteria to determine whether the system’s equilibrium was stable or unstable to pulse perturbations. Given the qualitative nature of the relationships described in the models, equations 2, 6 and 7 may have uncertain results due to the sum of positive and negative quantities with no specified magnitude. For example, equation 4 could have a negative result if the magnitude of the negative feedback terms was bigger than the magnitude of the positive ones. We built 10000 quantitative matrices, drawing values for the relationships between variables from random uniform distributions (*a*_*ij*_~*U*(0, 1)) keeping the same sign pattern as the original qualitative model. This allowed us to explore different combinations of the relative strengths of the links in the system. In other words, we simulated 10000 different quantitative systems that could exist in the real world and asked how many of these could be stable. We repeated the stability analysis on these matrices and the proportion of quantitative systems that met the stability criteria was used as a measure of the system’s potential for stability. Secondly, we conducted the sensitivity analysis (Hosack et al., 2009) of the stability criteria to identify which relationships in the system predominantly drive the scope for the system to achieve sustainability.

#### 2.2.2 Responses to press perturbations

For any loop model with *n* variables there are *n* possible points of entry for a press perturbation, one for each variable; a table of predictions can be built to show the direction of the response of each variable to a press perturbation acting on itself or on any other variable. The matrix of predictions is given by the adjoint of the system’s matrix. Each prediction represents the change in the equilibrium value (*) of a variable *X*_*j*_ due to a change in the parameter c that regulates the growth or the level of activity of a variable *X*_*i*_.

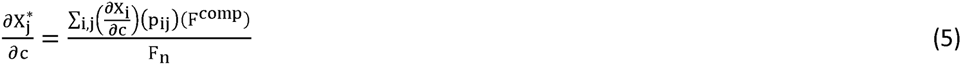

Each prediction is the result of the sum of all direct and indirect paths (*p*_*ij*_)) from the perturbed variable (*X*_*i*_)) to the response variable (*X*_*j*_)) multiplied by the direction of the change in the perturbed variable 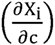 and by the feedback of their complementary subsystems (the subsystems of variables and links not included in a given path; F^comp^), divided by the overall feedback of the system (*F*_n_)).

Each element in the sum can be positive, negative or 0 and the magnitudes of the effects are not specified. Therefore, the prediction can have a certain degree of sign indeterminacy that can range from completely undetermined to uncertain predictions. Each prediction in the adjoint matrix can be weighted by the total number of cycles contributing to it, which is called the absolute feedback; the ratio between the absolute value of each element of the adjoint matrix and the corresponding value of absolute feedback gives the weighted-prediction matrix (Dambacher et al., 2002). Weighted predictions are a measure of uncertainty and range from 0 to 1; values near 0 represent predictions that are highly indeterminate, while values of 1 indicate that the prediction is completely reliable in terms of its sign. For each of the governance systems previously described, we used these predictions and their associated uncertainty to investigate how a tourism SES under different management scenarios will respond to a press perturbation.

The R package “LoopAnalyst” (Dinno, 2013; R Core Team, 2015) was used for conducting stability analysis and producing prediction tables, while sensitivity analysis was conducted in MATLAB (version 8.3.0.532, release 2014a, The MathWorks, Inc., Natick, Massachusetts, United States). R and MATLAB code are available as online supporting information (Appendices A & B respectively).

## 3. Results

### 3.1. Potential and conditions for stability to pulse perturbations

None of the systems was unconditionally stable (Table 1). The two users’ group ownership scenarios presented the highest potential for stability, followed by the open-access scenario; after a pulse perturbation, these systems have the highest chance to go back to their equilibrium state. A tourism SES under all the other scenarios has a good chance to be displaced from its equilibrium state after a small perturbation and either move to a different equilibrium, or present oscillatory instability (Levins, 1974; Puccia and Levins, 1985). Sensitivity analysis showed that the self-effect of E was consistently destabilising in all the scenarios (Appendix C). The environment needs to be at or close to its carrying capacity, where its rate of change is at its minimal. The positive loop between T and C was also crucial in determining stability in all the scenarios (Appendix C). In order for the system to be stable, infrastructures should not be a strong attractor for tourists and only a small proportion of tourists’ revenues should be reinvested in building new infrastructure.

**Table 1.** Potential for stability for all the models expressed as the proportion of quantitatively specified systems that met all stability criteria

| Model | Potential for<br>stability |
| --- | --- |
| Open access | 10.48% |
| Subsidies | 4.24% |
| Licensing | 7.98% |
| Access fee | 7.17% |
| User group ownership 1 | 21.42% |
| User group ownership 2 | 18.72% |
| Hybrid 1 | 5% |
| Hybrid 2 | 2% |

Some management strategies were destabilising for the system. In the subsidies scenario, the state intervention to subsidise the industry decreased potential for stability (Table C.2), while in the other two state ownership scenarios, the limiting strategies put in place by the government (licences and access fee) stabilised the system by creating negative feedback (Tables C.3 and C.4). However, the second stability criteria was usually sensitive to these negative links (Tables C.3 and C.4); these strategies tend to create long negative feedbacks that can potentially overwhelm short ones, thus decreasing potential for stability.

In both the users’ group ownership and the hybrid scenarios, the positive path from T to C through U, which represents the users’ alienation right, was crucial in determining stability (Tables C.5, C.6, C.7 and C.8). These links add positive feedback to the system, which tends to move the system away from its equilibrium state after a perturbation; however, they also create positive long feedbacks that counterbalance negative short ones, thus decreasing the probability of oscillatory instability of the system after a perturbation (Puccia and Levins, 1985).

The adaptive management strategy implemented by the government (state ownership and first hybrid scenarios; Tables C.2, C.3, C.4 and C.7), the users (second user group ownership scenario; Table C.6) or by the agency funded by the users (second hybrid scenario; Table C.8) to maintain environmental quality always increased the potential for stability.

### 3.2. Predictions and Triple Bottom Line sustainability

The open access scenario did not present any potential for TBL sustainability; most of the predictions (Table 2) showed negative responses of the variables to positive press perturbations to the system.

**Table 2.**
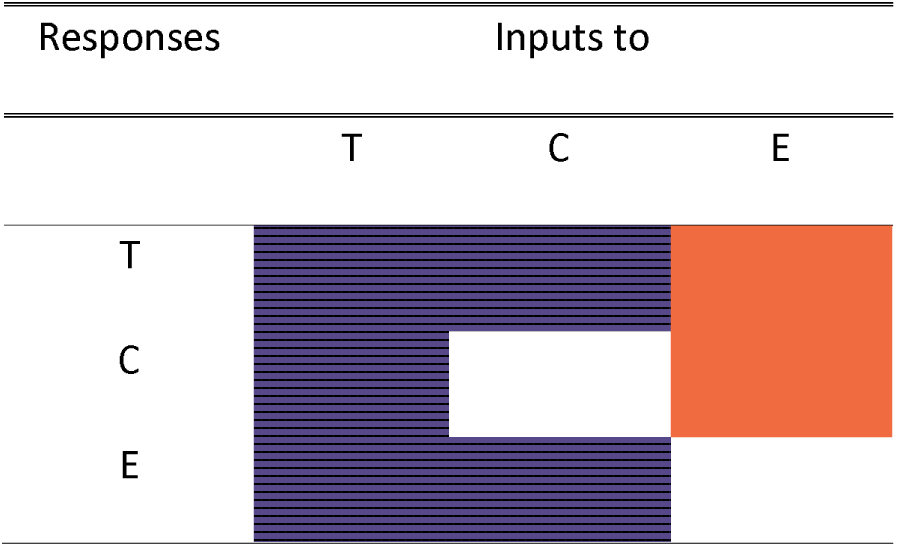
Predictions of responses to press perturbations for the open access scenario. In response to an increase in the column variable, the equilibrium value of the variable in the corresponding row either increased (orange), decreased (purple/striped), or we could not determine the response qualitatively (white). T: tourists; C: capital; E: environment

In the three state ownership scenarios, the environment did not respond to any press perturbation in the system, except for perturbations to S (Table 3). State intervention acted as a buffer of the environment, absorbing all the press perturbations that enter the system (Puccia and Levins, 1985). This result highlights the importance of an adaptive strategy to natural resource management. Only the subsidies scenario was not compatible with the concept of TBL sustainability, while a licencing scenario offered scope for the industry to grow sustainably (Table 3). However, in this scenario, it is uncertain how the capital will respond to an increase in the number of tourists. There are two ways T can influence C (Fig. 1b, blue dash-dotted line): the direct effect is positive, while the indirect path is negative (an increase in T has a negative effect on E, which will stimulate S to reduce C). When the direct positive feedback cycle is stronger than the indirect negative one, the response of the capital will be positive and TBL sustainability achievable. This condition is opposite to the conditions for stability; among the simulated quantitative systems, most of the stable ones showed a negative response of C to increases in T, even though positive responses were also possible (Fig.D.1). This indicates that TBL sustainability is possible, but very difficult to achieve in a licencing scenario. The same was true for the response of T to an increase in C in the “access fee” scenario (Table 3). An expansion of infrastructure might degrade the environment, which will lead the government to restrict the access to the area from the tourists by increasing the access fee. If the tourists are strongly attracted by the infrastructure and the response of the government is not too strong, the number of tourists might not decline, but the system will be unstable to pulse perturbations.

**Table 3.**
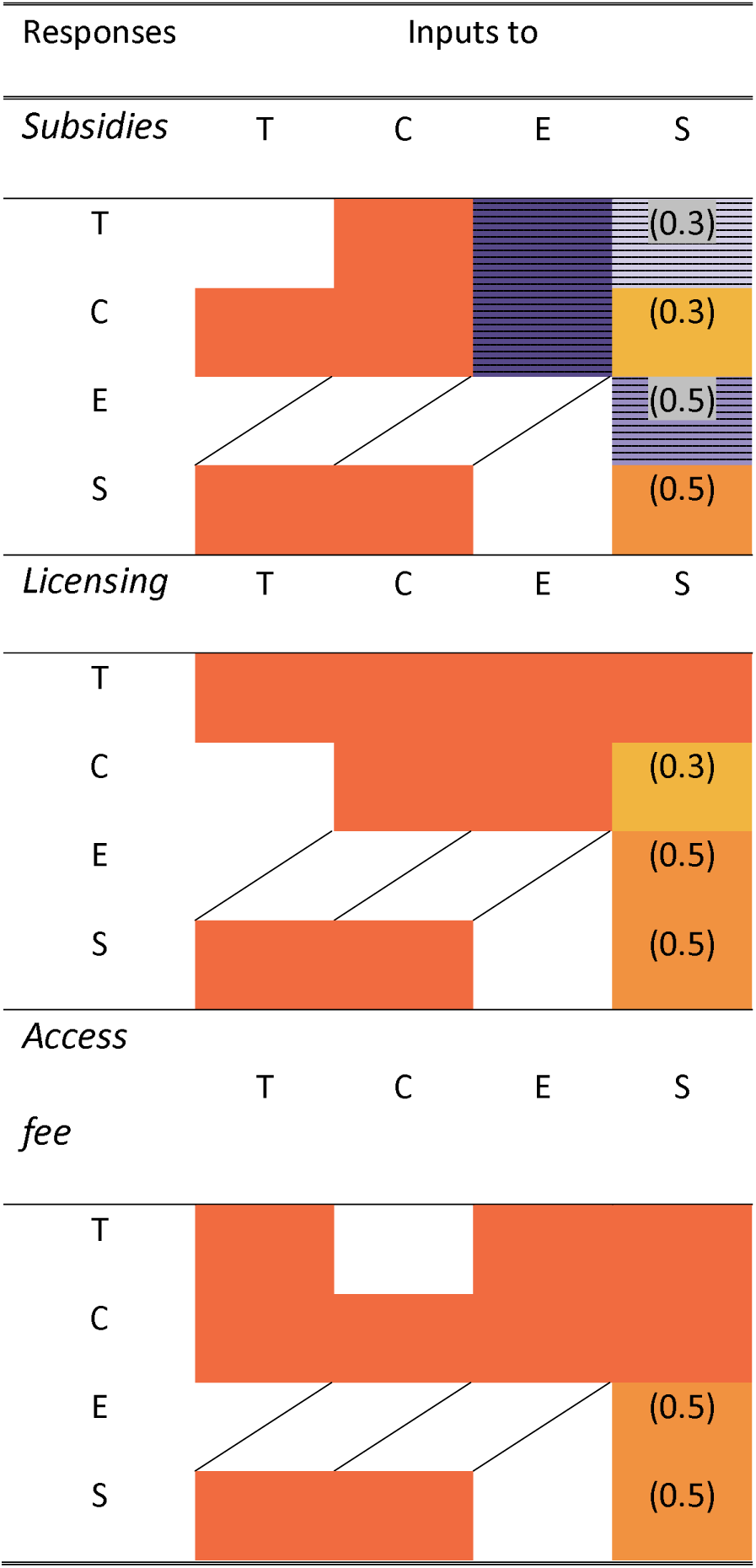
Predictions of responses to press perturbations for state ownership scenarios. The equilibrium value of the variable in the corresponding row either increased (orange), decreased (purple/striped), or was not affected (white, crossed) in response to an increase in the column variable. Some responses could not be determined qualitatively (white) and for ambiguous responses (shaded orange or purple/striped) we provide values of weighted feedback. Values of 0.5 give a sign determination that exceeds 90% (Dambacher, Li & Rossignol 2003). T: tourists; C: capital; E: environment; S: state intervention

Group property rights regimes, which showed higher resilience to pulse perturbations, had no potential for TBL sustainability (Table 4). In the first scenario, there is a high number of negative responses to positive inputs to the system. The second scenario showed a very high degree of indeterminacy (values of weighted feedback < 0.5 (Dambacher et al., 2003)) and some negative responses, which means that TBL sustainability is not likely to be achieved.

**Table 4.**
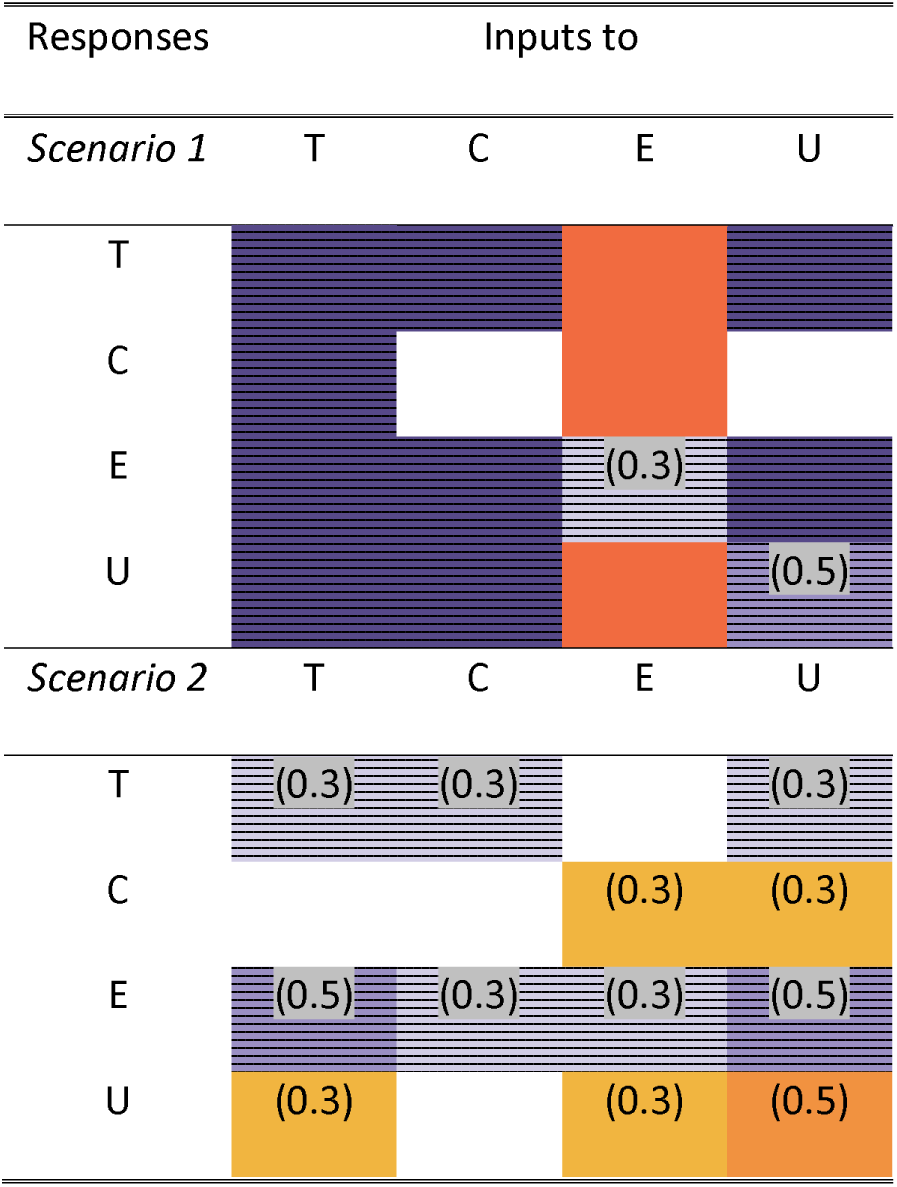
Predictions of responses to press perturbations for users’ group ownership scenarios. The equilibrium value of the variable in the corresponding row either increased (orange) or decreased (purple/striped) in response to an increase in the column variable. Some responses could not be determined qualitatively (white) and for ambiguous responses (shaded orange or purple/striped) we provide values of weighted feedback. Values of 0.5 give a sign determination that exceeds 90% (Dambacher, Li & Rossignol 2003). T: tourists; C: capital; E: environment; U: users

In contrast, the two hybrid scenarios had predictions compatible with the concept of TBL sustainability. There was only one negative response in the first scenario (Table 5): following an increase in state intervention, the environment could degrade. This counter-intuitive response was uncertain (weighted feedback = 0.3; Table 5). Moreover, the conditions for this response to be positive were the same as the conditions for stability and this response was always positive in quantitatively stable systems (Fig.D.2a). The undetermined predictions of the response of users to an increase in the number of users could potentially be a problem for social justice. If the negative self-loops of T, C and U are weaker than the positive loop between T and C, then the number of users decreases, with a potential for monopolisation. This condition is never satisfied in stable systems, so the response of U to inputs to U is always positive in stable systems and TBL sustainability achieved (Fig.D.2b). The second scenario showed more uncertainty (Table 5). An increase in tourism structures gave an undetermined response of the capital itself. Conditions for this response to be positive were the same as conditions for stability and the capital always responded positively to positive inputs to the capital in quantitative stable systems (Fig.D.3a). The same was true for the response of the environment and users to an increase in users (Fig. D.3b & c) and the response of the environment to an increase in the management effort of the agency (Fig.D.3c). These responses were always positive in quantitative stable systems. Therefore, this scenario has the potential to be sustainable according to the TBL sustainability concept in presence of conditions for stability to pulse perturbations.

**Table 5.**
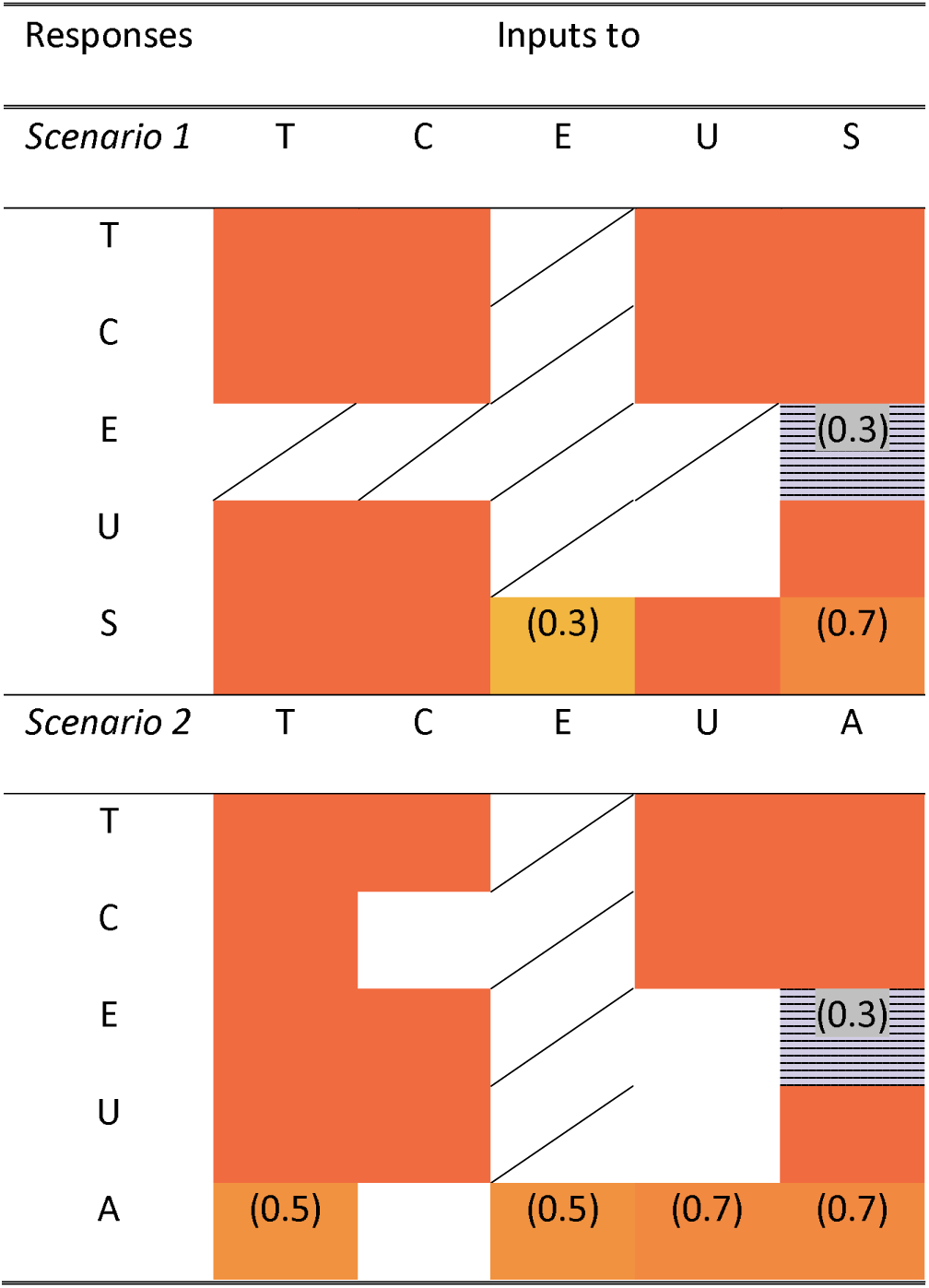
Predictions of responses to press perturbations for hybrid scenarios. The equilibrium value of the variable in the corresponding row either increased (orange), decreased (purple/striped), or was not affected (white, crossed) in response to an increase in the column variable. Some responses could not be determined qualitatively (white) and for ambiguous responses (shaded orange or purple/striped) we provide values of weighted feedback. Values of 0.5 give a sign determination that exceeds 90% (Dambacher, Li & Rossignol 2003). T: tourists, C: capital, E: environment, U: users, A: external agency

## 4. Discussion

A qualitative approach to SESs modelling provided a way to test alternative governance structures and assess whether they would influence the sustainability of nature-based tourism. SESs are subjected to press-pulse dynamics (Collins et al., 2011) and in order to predict the outcomes of different management strategies we need to investigate their responses to both press and pulse perturbations. In order to be sustainable, a SES needs to be resilient to pulse perturbations and, in presence of a press change, such as economic growth, the system needs to maintain TBL sustainability.

Here we showed that in instances when nature-based tourism systems can be considered exploiting a common good (Pirotta and Lusseau, 2015) then they are most likely to be unstable, regardless of the management strategy adopted. We confirm the results from previous studies (Casagrandi and Rinaldi, 2002) that showed that in open access tourism SESs, sustainability is often at risk because small perturbations can have dramatic effects. We generalise this finding to the main governance structures available for common goods. A small pulse perturbation can potentially drive the system away from its equilibrium and either move it to a new, unknown, state or cause oscillations. This is an unfavourable property, because it makes the SES, on which many livelihoods depend on, unreliable and vulnerable to any disturbance.

However, all the systems tested in this study had some potential for local stability to pulse perturbations, which means that by modifying some variables in the system, a reliable and resilient tourism SES is achievable. First, it is important to understand what attracts tourists to a site, because a strong demand for tourism infrastructures is very likely to lead to instability. Secondly, in order to maintain stability, a higher proportion of the tourism revenues should be invested in renewing old infrastructures, instead of investing in new ones. Thirdly, the exploited common resource needs to be maintained in a state where its rate of change is minimal; for example, keeping a wildlife population close to its carrying capacity. The self-enhancing effect that results from the resource being exploited to the point that it is far from its ‘pristine’ abundance/density strongly affects the resilience of the system. This result agrees with many studies that have discouraged the application of the Maximum Sustainable Yield (MSY) concept in the management of harvested populations and ecosystems, on the basis that it would lead to extinction of some species instead of guaranteeing a sustainable use (Geček and Legović, 2012).

Some management scenarios exhibited higher potential for stability than others. However, this potential for stability to pulse perturbations did not always correspond to the potential for sustainable development of the industry. For instance, open access and user group ownership scenarios showed the highest potential for stability (Table 1), but they had no potential to achieve TBL sustainability (Tables 2 & 4). In open access commons, overexploitation of the resource happens because the perceived benefits of overuse are always higher than the perceived losses (Hardin, 1968) and users have no incentives to invest in the resource or conserve it for the future (Acheson, 2006). Therefore, without any regulation, a CPR is doomed to degradation and human activities to failure. Local knowledge of user groups can confer more resilience to user-managed SESs (Berkes et al., 2003), but it does not guarantee sustainable growth (Table 4) (Ostrom, 1990).

State ownership scenarios were very sensitive to pulse perturbations (Table 1), but two of them offered a better outlook for TBL sustainability. The licensing and the access fee scenario could potentially lead to a stable system that has scope for sustainable growth, but this outcome was possible only in a very narrow range of parameter space. Conditions for stability (Appendix C) contrasted with conditions for TBL sustainability (Table 3), therefore only a few quantitative stable systems had scope for sustainable growth (Fig. D.1). A high rate of investment of tourists’ revenues into infrastructures would assure an increase in the tourism capital following an increased affluence of tourists, which is a good prediction in terms of economic viability of the industry and employment opportunities. However, this condition would make the system unstable, so vulnerable to any small perturbation. A negative response of the capital to an increase in the number of tourists could be a problem in terms of social justice; a high number of tourists could result in a decrease in environmental quality, for example due to littering, pollution or excessive disturbance on the wildlife. This would cause the government to reduce the number of licenses available to the tour operators, which would exclude some of them from the industry. These results indicate that it might be very difficult to find a balance between all these conditions and design effective rules, for example deciding how many licences should be issued or the price of the access fee (Acheson, 2006). Since perfectly designed management strategies are rarely achieved, sustainable development in SESs governed by centralised institutions is possible only by trading off some of the system’s robustness to pulse perturbations.

Nonetheless, locally user-defined market-based strategies can fail too. The alienation right, represented in our models by the positive links between T, U and C (Fig. 1c & d), was highly destabilising for the system (Appendix C). Introducing positive feedback into the system contributed to destabilisation. Previous studies have suggested that simple strategies, where the market or the government alone have complete control over the resource, often fail and that a combination of different institutional arrangements creates better conditions for sustainable governance (Dietz et al., 2003; Meinzen-Dick, 2007; Pirotta and Lusseau, 2015). We showed that in a management regime where property rights and responsibilities are shared between the users and a third party there is a good potential for sustainable development of the industry (Table 5). In these strategies, users still retain their rights of access and use, exclusion and alienation, but the management of the resource is left to the government (Fig. 1d, orange dotted lines) or an external agency funded by the users (Fig. 1d, green dashed lines). These scenarios are very sensitive to pulse perturbations, which means that there is a narrow range of parameter space where these systems can be sustainable and grow. However, sustainability is achievable by careful tuning of the relative strengths of the relationships in these systems. A nature-based tourism SES where there is a low demand for tourism infrastructure, a low reinvestment rate of tourists’ revenues into new infrastructure and a healthy environment could be sustainable in presence of press-pulse dynamics under a hybrid governance regime.

Using an integrated qualitative approach that takes into account economic profitability, environmental quality and a form of social justice, we have identified which management regimes have the highest potential for sustainability, and the conditions necessary for them to achieve it. We agree with Ostrom’s Law that one-size-fits-all solutions fail in most real situations. This happens because SESs are highly unstable and sustainability is only possible in very narrow regions of parameter space. We have showed that some management strategies have higher potential for sustainability than others, but each strategy must be carefully tuned to each particular situation. Although our models of a tourism-based system are extremely simplified, they are representative of all the main components of Ostrom’s conceptual map (Ostrom, 2007): resource system, users, governance system and their interactions. More detailed SESs can now be explored, by unpacking these highest-tier conceptual variables (Ostrom, 2007). We propose that this qualitative approach can be a powerful diagnostic tool to identify variables and their combinations that affect sustainability of different governance systems in common-pool resource management.

## Acknowledgments

This work was funded by the University of Aberdeen and Scottish Natural Heritage (SNH) and their support is gratefully acknowledged. We thank MASTS (the Marine Alliance for Science and Technology for Scotland) for their role in funding this work and B. Leyshon and F. Manson (SNH) for fruitful discussion.

